# Halo: a pretrained model for whole-cell segmentation from nuclei images in spatial transcriptomics

**DOI:** 10.64898/2026.04.02.716237

**Authors:** Xingyuan Zhang, Haotian Zhuang, Zhicheng Ji

## Abstract

Spatial transcriptomics enables measurement of gene expression while preserving spatial organization within tissues. Accurate reconstruction of single-cell transcriptomes requires precise whole-cell segmentation, yet many spatial transcriptomics experiments provide only nuclear staining images, making reliable inference of cell boundaries difficult. Here we introduce Halo, a pretrained segmentation model that reconstructs whole-cell boundaries by integrating nuclear morphology with the spatial distribution of RNA transcripts. Halo converts transcript coordinates into molecular density maps that are processed jointly with DAPI images using a Cellpose-SAM segmentation architecture. Unlike existing approaches that require dataset-specific training, Halo is pretrained on multimodal Xenium data from 12 tissue types and can be directly applied to new datasets without additional training. Across diverse tissues, Halo substantially outperforms the widely used nuclear expansion strategy, achieving higher agreement with ground-truth cell boundaries and more accurate RNA-to-cell assignment. Improved segmentation leads to more reliable cell type identification and more accurate estimation of cell morphological features. By providing a pretrained, generalizable model for whole-cell reconstruction, Halo enables scalable and reproducible cell segmentation for image-based spatial transcriptomics.

## Introduction

Spatial transcriptomics (ST) is a rapidly evolving technology that enables the measurement of gene expression while preserving the spatial organization of cells within tissues, providing critical insights into tissue architecture, cell–cell interactions, and spatially organized biological processes^1–4^. Broadly, ST platforms can be categorized into spot-based^5,6^ and image-based approaches^7,8^. Spot-based methods profile transcripts captured within predefined spatial locations, or spots, each of which typically contains RNA from multiple neighboring cells, resulting in limited spatial resolution. In contrast, image-based methods detect transcripts directly within tissue sections using imaging and in situ hybridization or sequencing strategies, achieving single-cell or even subcellular resolution. Owing to this finer spatial granularity, image-based approaches enable more precise characterization of cellular heterogeneity and spatial gene expression patterns, and are particularly powerful for resolving complex tissue microenvironments.

In addition to providing spatial locations of RNA transcripts, image-based ST technologies also generate complementary fluorescence imaging data. For example, the widely used 10x Xenium platform offers two imaging configurations. The first includes DAPI staining for imaging cell nuclei, while the second, multimodal configuration additionally captures signals from cell boundaries, intracellular RNA, and other molecular markers alongside DAPI staining. This rich imaging information makes it possible to infer cell locations and boundaries through a computational procedure known as cell segmentation^9–12^. Cell segmentation is a critical step for assigning transcripts to individual cells and constructing single-cell gene expression matrices, which serve as the basis for most downstream analyses, including cell type annotation^13,14^, spatial domain detection^15^, and identification of spatially variable genes^16,17^.

Ideally, cell segmentation aims to reconstruct whole-cell boundaries rather than only nuclear boundaries, because transcripts located in the cytoplasm carry important information about cell identity^18^. However, accurate whole-cell segmentation remains challenging when only DAPI-stained images are available, as these images label only nuclei and provide little direct information about the surrounding cytoplasm or cell membrane. Even when multimodal imaging is available, which often requires substantially greater financial and experimental resources, tens of thousands of cells may still lack reliable signals from additional imaging channels and therefore rely primarily on nuclei staining to infer whole-cell boundaries (Figure 1a). In such settings, existing image-based segmentation methods^9–11^ can reliably delineate nuclei but generally cannot reconstruct accurate whole-cell boundaries.

**Figure 1.**
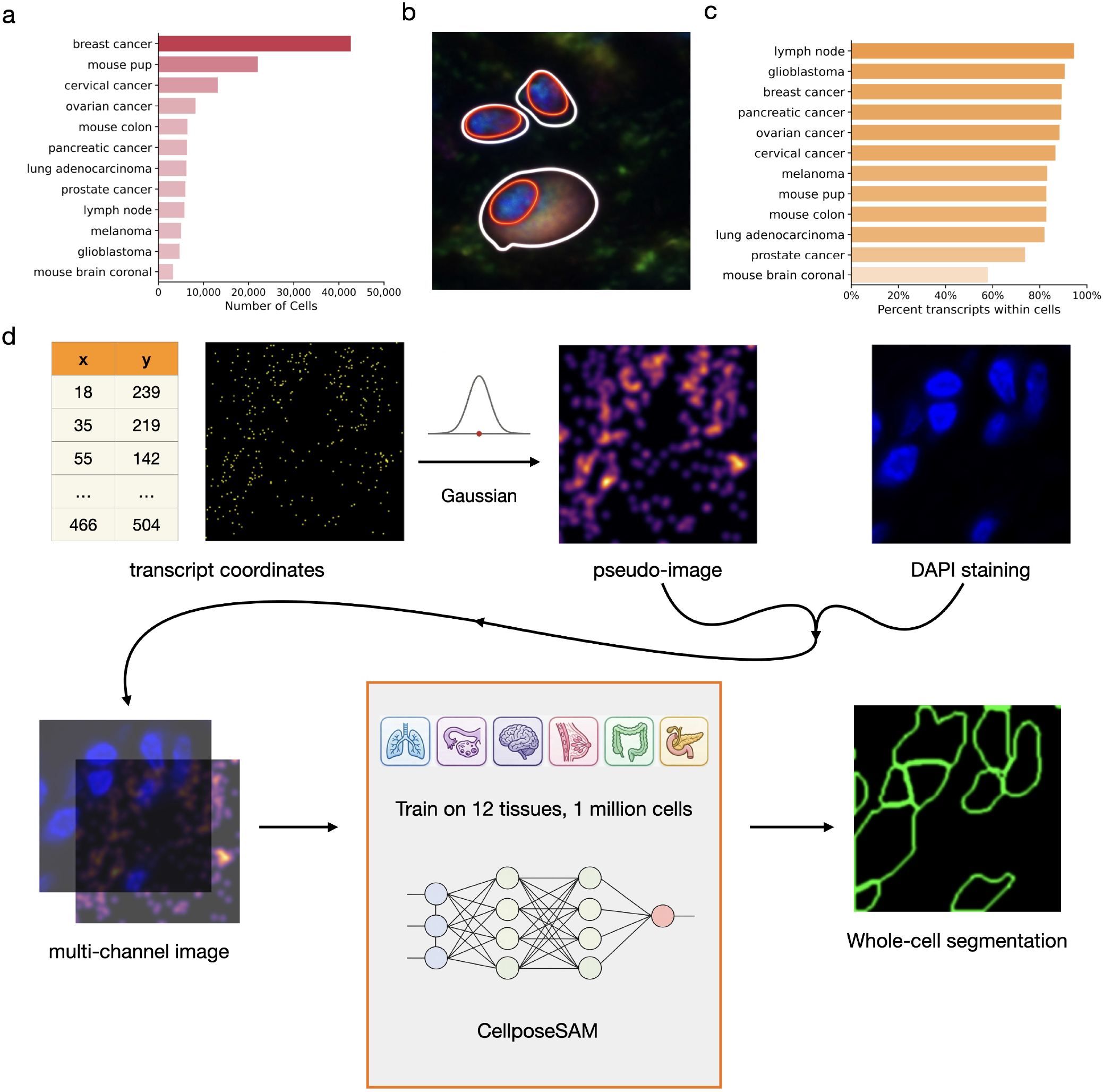
**a**, Number of cells segmented based on nuclei staining across Xenium datasets with multimodal imaging. **b**, Whole-cell boundaries (white contours) and nuclear boundaries (red contours) for three representative cells. **c**, Percentage of RNA transcripts within cell boundaries across Xenium datasets. **d**, Overview of Halo.

A widely adopted strategy, implemented in the 10x Space Ranger analysis pipeline, addresses this limitation by first segmenting nuclei from DAPI images and then expanding each nuclear mask by a fixed number of pixels to approximate the corresponding whole-cell boundary. However, this approach has several major limitations (Figure 1b). First, whole-cell morphology can differ substantially from nuclear morphology. Second, the distance between the nucleus and the cell boundary varies widely across cells. Third, the nucleus is not always centrally located within a cell. As a result, uniformly enlarging nuclei by a fixed number of pixels often fails to accurately capture true cell boundaries and can lead to inaccurate assignment of transcripts to cells.

Several recent computational methods have attempted to improve segmentation accuracy by integrating RNA spatial coordinates with imaging information^19,20^. These approaches demonstrate that the spatial distribution of transcripts contains useful information for delineating cellular boundaries. However, existing methods typically require dataset-specific model training and annotated segmentation masks, which limits their applicability across datasets and tissue types. In practice, many spatial transcriptomics datasets lack high-quality ground-truth segmentation masks, making it difficult to apply such approaches broadly.

To address this limitation, we developed Halo, a pretrained segmentation model that reconstructs whole-cell boundaries by integrating nuclear morphology with the spatial distribution of RNA transcripts. Halo leverages the observation that most transcripts reside within cells and therefore collectively encode information about cellular extent (Figure 1c). To integrate transcript spatial information with nuclear imaging, Halo converts RNA coordinates into molecular density maps that can be processed jointly with DAPI images within a modern segmentation architecture. The model is trained on multimodal Xenium datasets spanning 12 tissue types, enabling it to learn generalizable relationships between nuclear morphology, transcript spatial patterns, and whole-cell structure. As a result, Halo can be directly applied to new spatial transcriptomics datasets without requiring additional model training or annotated segmentation masks.

We demonstrate that Halo accurately reconstructs whole-cell boundaries across diverse tissue types and substantially outperforms the commonly used nuclear expansion strategy. Improved segmentation leads to more accurate RNA-to-cell assignment, better cell type identification, and more reliable estimation of cell morphological features. By providing a pretrained and generalizable model for whole-cell reconstruction, Halo enables scalable and reproducible cell segmentation for image-based spatial transcriptomics analyses.

## Results

### Halo overview

Halo predicts whole-cell boundaries by jointly leveraging RNA spatial coordinates and DAPI nuclear staining images. To integrate these heterogeneous modalities, it first converts RNA spatial location data into pseudoimages, transforming molecular information into an image-compatible representation that can be processed alongside nuclei images within a unified framework. A Cellpose-SAM model^21^, a recent foundation model for biological segmentation, is then trained on multi-channel inputs comprising the RNA pseudoimage and the DAPI image to infer whole-cell boundaries. Halo is trained using multimodally stained samples generated on the 10x Xenium platform across 12 distinct tissue types (Figure 1d), enabling robust performance and broad applicability across diverse tissue contexts. Halo can be directly applied to a new dataset using RNA spatial coordinates and DAPI nuclear staining images as inputs.

### Halo improves whole-cell segmentation and transcript assignment

We applied Halo to image tiles in the test set that were not used during Halo’s training. As a competing approach, we also applied the nuclear expansion method. Figure 2 compares the ground-truth whole-cell segmentation with the segmentation results obtained by Halo and nuclear expansion in four representative regions from different tissues. The segmentation produced by Halo is highly consistent with the ground truth. In contrast, the nuclear expansion results deviate from the ground truth in most cases, exhibiting two major issues. First, nuclear expansion typically produces round or elliptical boundaries that fail to capture the more complex cell boundary contours observed in the ground truth. Second, the boundaries generated by nuclear expansion are generally larger than those in the ground truth.

**Figure 2.**
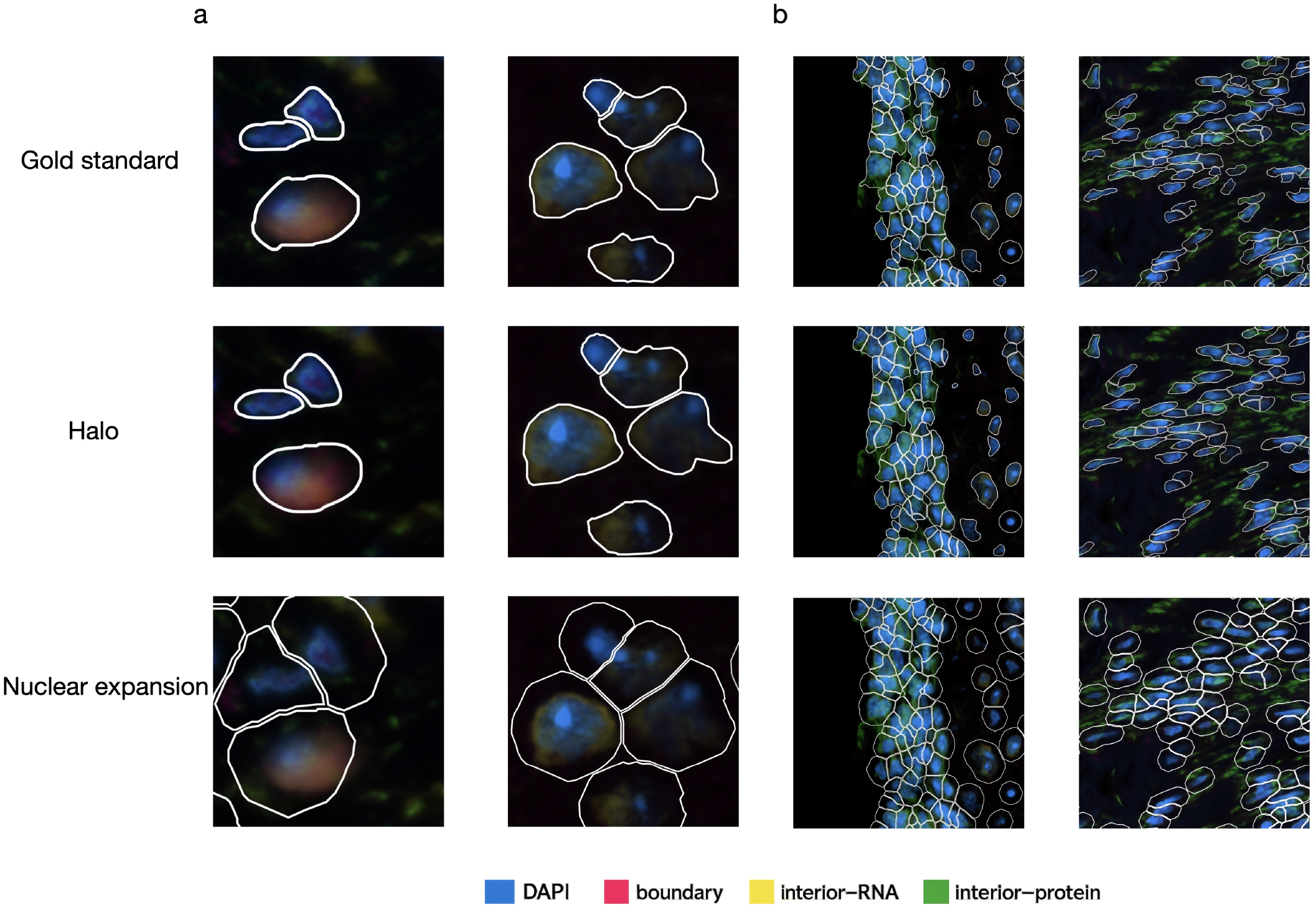
Whole-cell segmentations (white boundaries) of representative cells for ground truth, Halo, and nuclear expansion. **a**, Zoomed-in examples, left, human pancreatic cancer, right, mouse brain coronal. **b**, Zoomed-out examples, left, human glioblastoma, right, human melanoma.

We then systematically evaluated the performance of Halo and nuclear expansion across different tissue types, using image Intersection over Union (IoU) to quantify the agreement between whole-cell masks generated by Halo or nuclear expansion and the ground truth (Figure 3). Halo achieves higher image IoU values than nuclear expansion for every tissue type. Across all tissues, Halo reaches a median IoU of approximately 0.7, which is about 0.15 higher than that of nuclear expansion. These results indicate that the whole-cell boundaries produced by Halo are highly consistent with the ground truth and significantly more accurate than those obtained using nuclear expansion.

**Figure 3.**
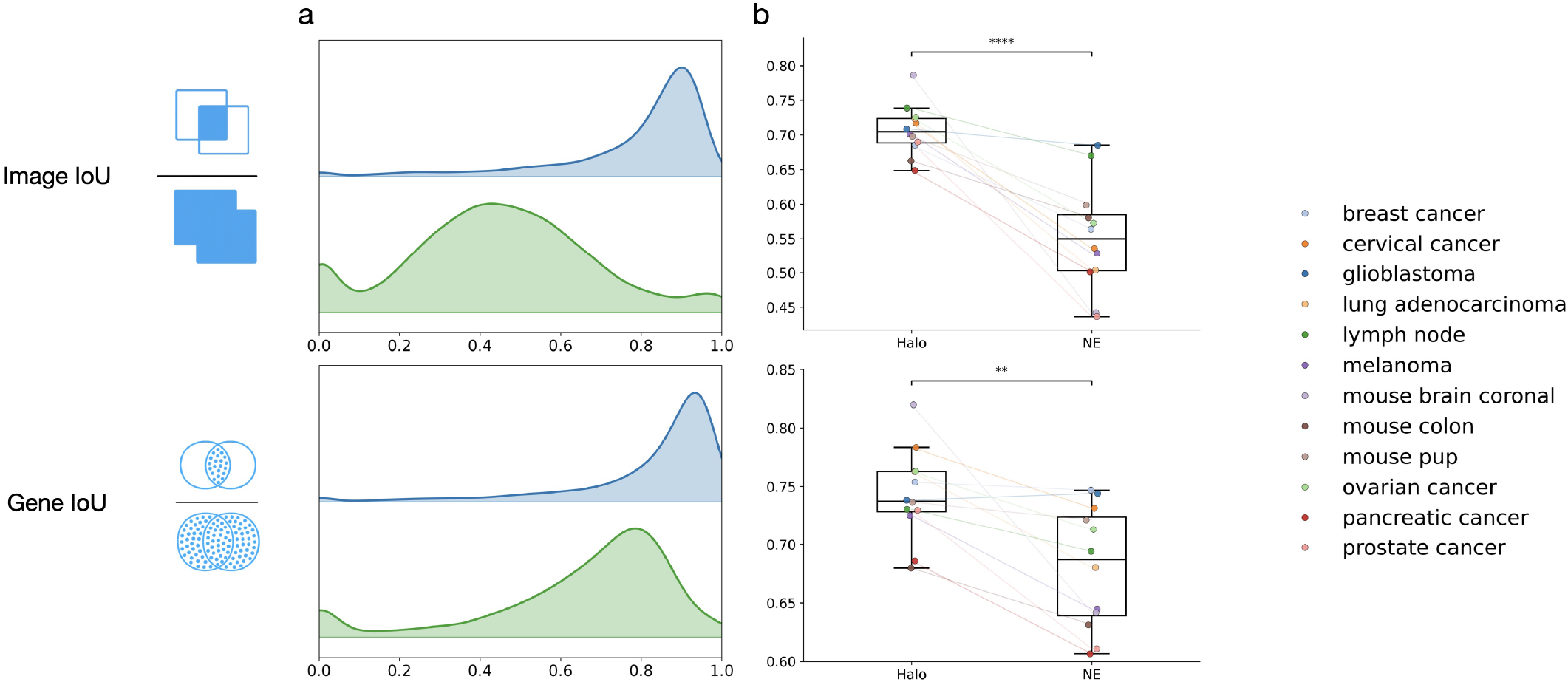
Evaluation of Image IoU and Gene IoU comparing Halo with nuclear expansion. **a**, Distributions of Image IoU (top) and Gene IoU (bottom) across cells in a representative mouse brain coronal dataset. **b**, Mean Image IoU (top) and mean Gene IoU (bottom) across datasets. **** indicates p-value < 0.0001, and ** indicates p-value < 0.01.

Once whole-cell boundaries are obtained, an RNA transcript can be assigned to a cell if its spatial location falls within the cell boundary. To quantitatively evaluate this step, we devised gene IoU, a metric analogous to the image IoU described above, to assess the agreement between RNA–cell assignments derived from Halo or nuclear expansion and those from the ground truth (Figure 3). Halo again achieves higher gene IoU values than nuclear expansion in nearly every tissue type. Across tissues, Halo attains an overall gene IoU of nearly 0.75, which is significantly higher than that of nuclear expansion. These results indicate that the RNA–cell assignments produced by Halo are accurate and markedly better than those obtained using nuclear expansion.

### Halo improves cell type identification

Once RNA transcripts are assigned to cells and a single-cell count matrix is generated, a key task is to identify the cell type of each cell based on its transcriptomic profile. The accuracy of cell type identification directly affects several downstream analyses that rely on cell type information, such as cellular neighborhood and interaction analysis^22,23^, and the identification of cell-type-specific spatially variable genes^16,24^.

We applied the same cell type annotation pipeline to the gene expression count matrices derived from the gold-standard whole-cell boundaries, as well as from the boundaries identified by Halo and nuclear expansion. Figure 4 shows the cell type identification results produced by these methods in several example tissue regions. While the cell type annotations based on Halo largely agree with those based on the ground truth, the annotations derived from nuclear expansion exhibit clear discrepancies in several cases. For example, nuclear expansion can mistakenly annotate T cells as cancer cells, potentially biasing analyses of cancer–immune interactions.

**Figure 4.**
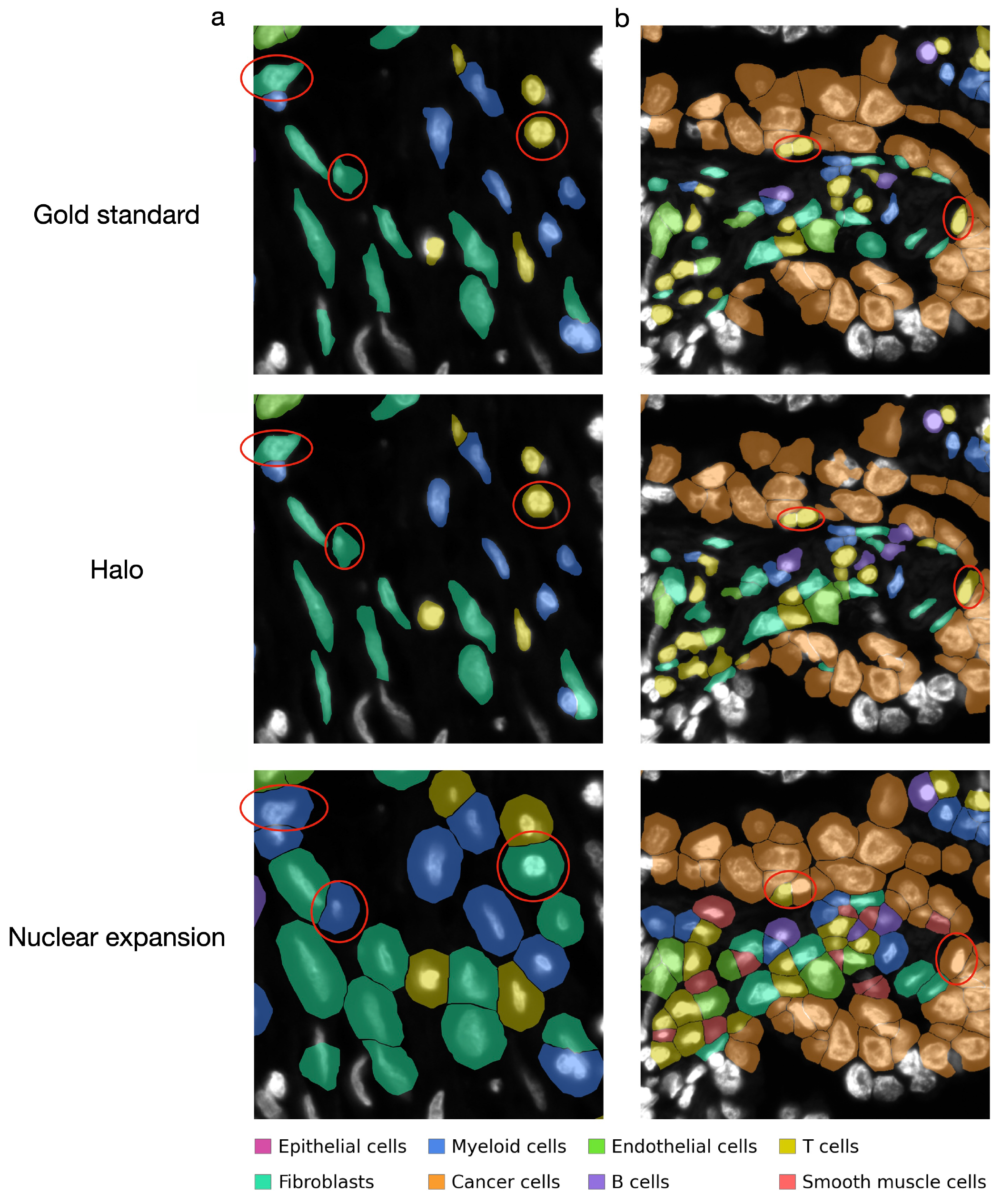
Examples of cell type annotation based on the gold standard, Halo, and nuclear expansion in human pancreatic cancer (**a**) and human lung adenocarcinoma (**b**). Cells incorrectly labeled by nuclear expansion are highlighted with red circles.

We further quantitatively compared the cell type identification results derived from Halo and nuclear expansion with those from the ground truth using metrics including the adjusted Rand index (ARI), adjusted mutual information (AMI), homogeneity, and completeness (Figure 5). Halo achieves better clustering metrics than nuclear expansion in nearly all tissues and shows significantly improved clustering performance overall across tissues. These results suggest that Halo enables more accurate cell type identification.

**Figure 5.**
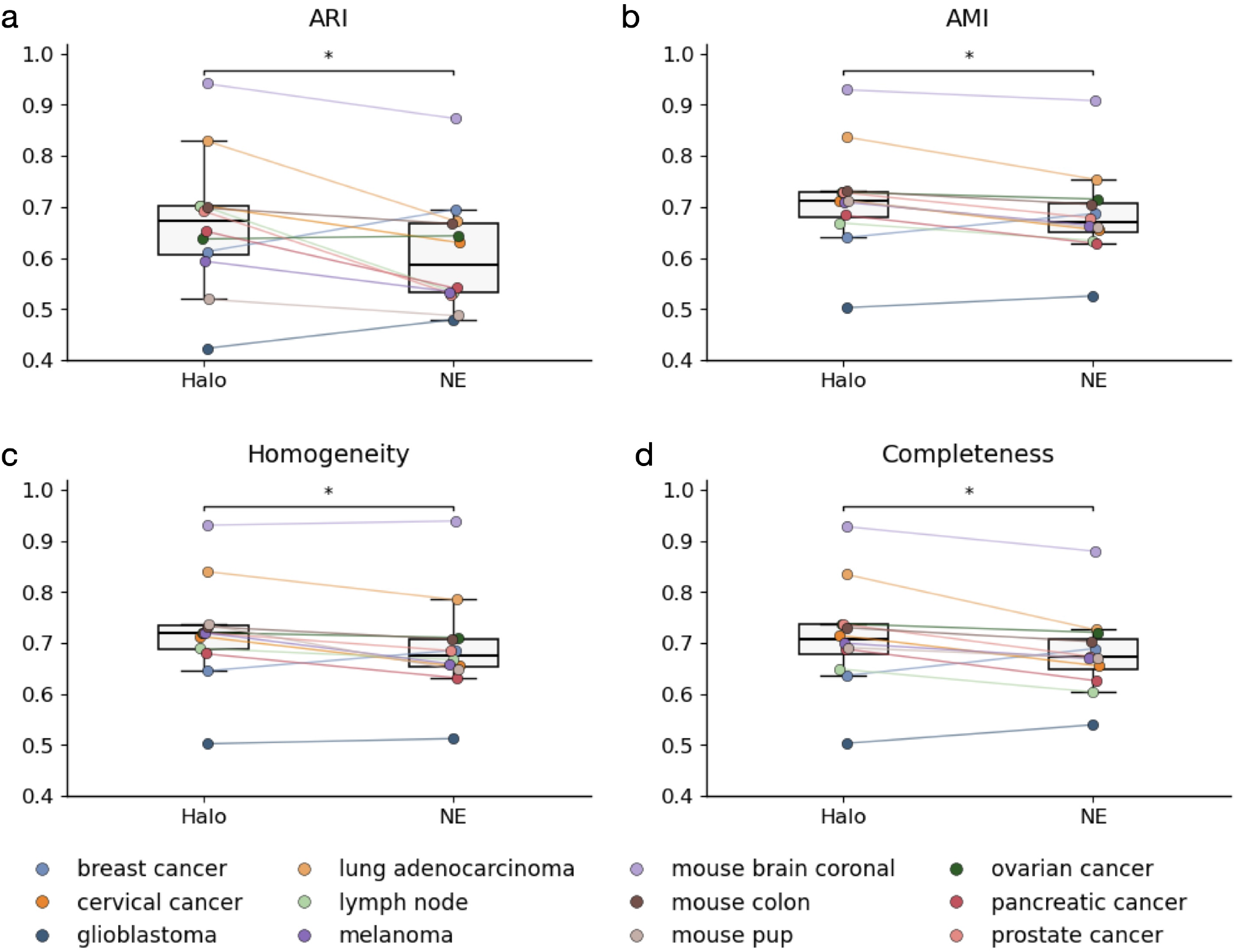
Evaluation of clustering performance for Halo and nuclear expansion. **a**, ARI. **b**, AMI. **c**, Homogeneity; **d**, Completeness. * indicates p-value < 0.05.

### Halo more accurately captures cell morphological features

By jointly profiling the morphological and transcriptomic features of the same cell, spatial transcriptomics enables the study of cell morphology in a cell-type-specific manner, which is important for understanding cellular states, disease progression, and tissue organization^25–27^. To evaluate whether the whole-cell segmentation produced by Halo can be used for morphological analysis, we calculated several morphological features, including cell size, aspect ratio, and roundness, using ground-truth whole-cell boundaries as well as boundaries derived from Halo or nuclear expansion (Figure 6). The morphological features obtained using Halo are highly consistent with those from the ground truth and reflect known biological characteristics. For example, lymphocytes such as T cells and B cells exhibit relatively small areas and high roundness, reflecting their compact and spherical morphology, whereas fibroblasts and smooth muscle cells show larger aspect ratios and lower roundness due to their elongated, spindle-like shapes. Epithelial cells tend to have larger cell areas, consistent with their sheet-like organization in tissues. These patterns agree with well-established observations from quantitative cell morphology studies, indicating that the extracted features capture biologically meaningful differences between cell types^28–30^. In contrast, nuclear expansion substantially distorts morphological features. For example, there is much less variation in aspect ratio and roundness across cell types, likely because the inferred whole-cell boundaries are circular or elliptical in most cases (Figure 2).

**Figure 6.**
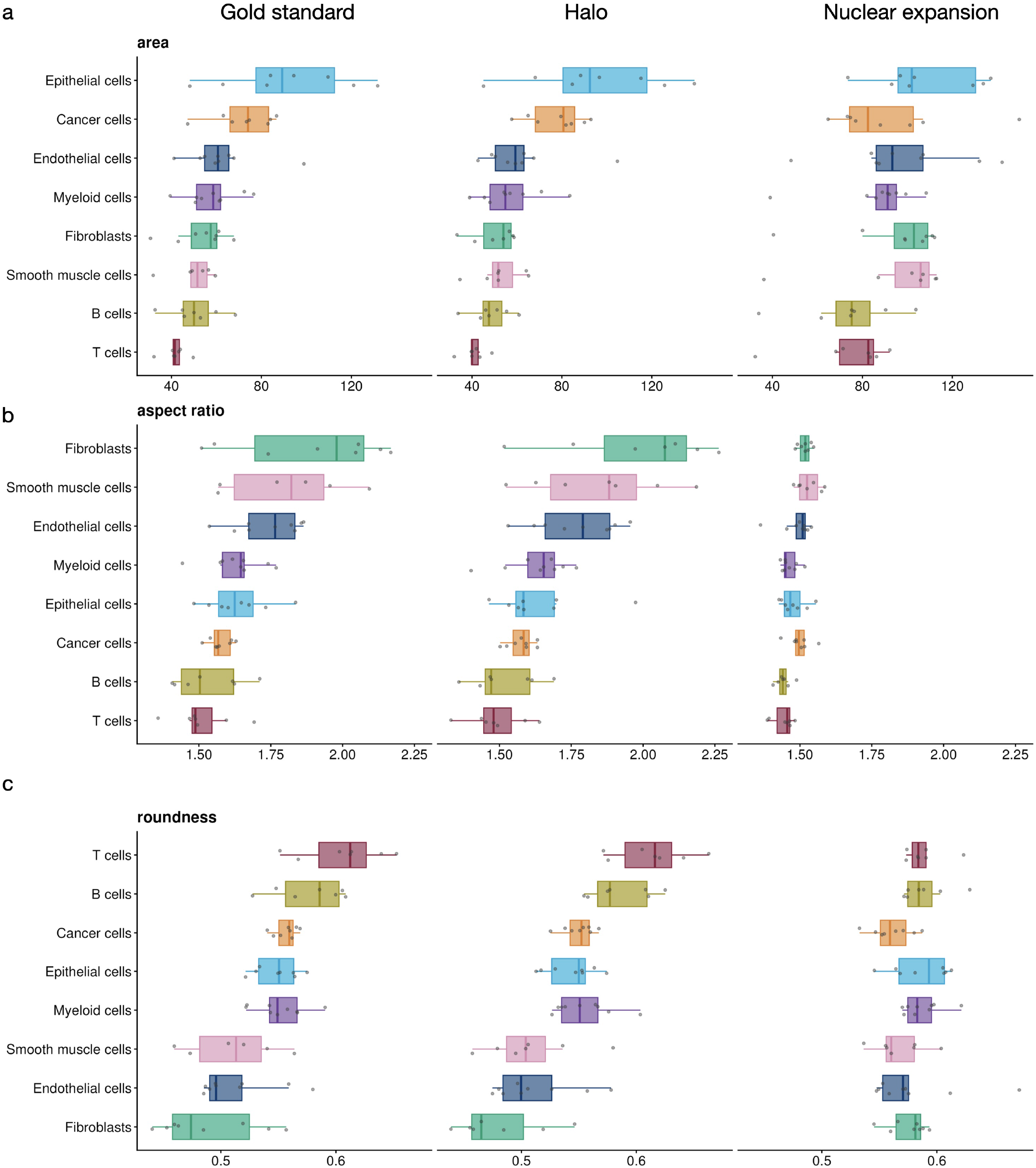
Distributions of morphological features across cell types for different segmentation methods. **a**, Cell area. **b**, Aspect ratio. **c**, Roundness.

Finally, we tested whether morphological features can be used to separate or predict different cell types. We first projected the morphological features calculated for each cell type within each tissue into a low-dimensional space using Uniform Manifold Approximation and Projection (UMAP) (Figure 7a). Similar to the ground truth, morphological features derived from Halo lead to clear separation of different cell types, while the same cell types from different tissues remain close to one another. In contrast, when morphological features derived from nuclear expansion are used, different cell types appear largely mixed in the UMAP embedding. We further evaluated cell type prediction accuracy in a cross-validation study using only morphological features as predictors. Halo achieves significantly higher prediction accuracy than nuclear expansion in nearly all scenarios. These results suggest that morphological features derived from Halo more effectively separate and predict different cell types across tissues.

**Figure 7.**
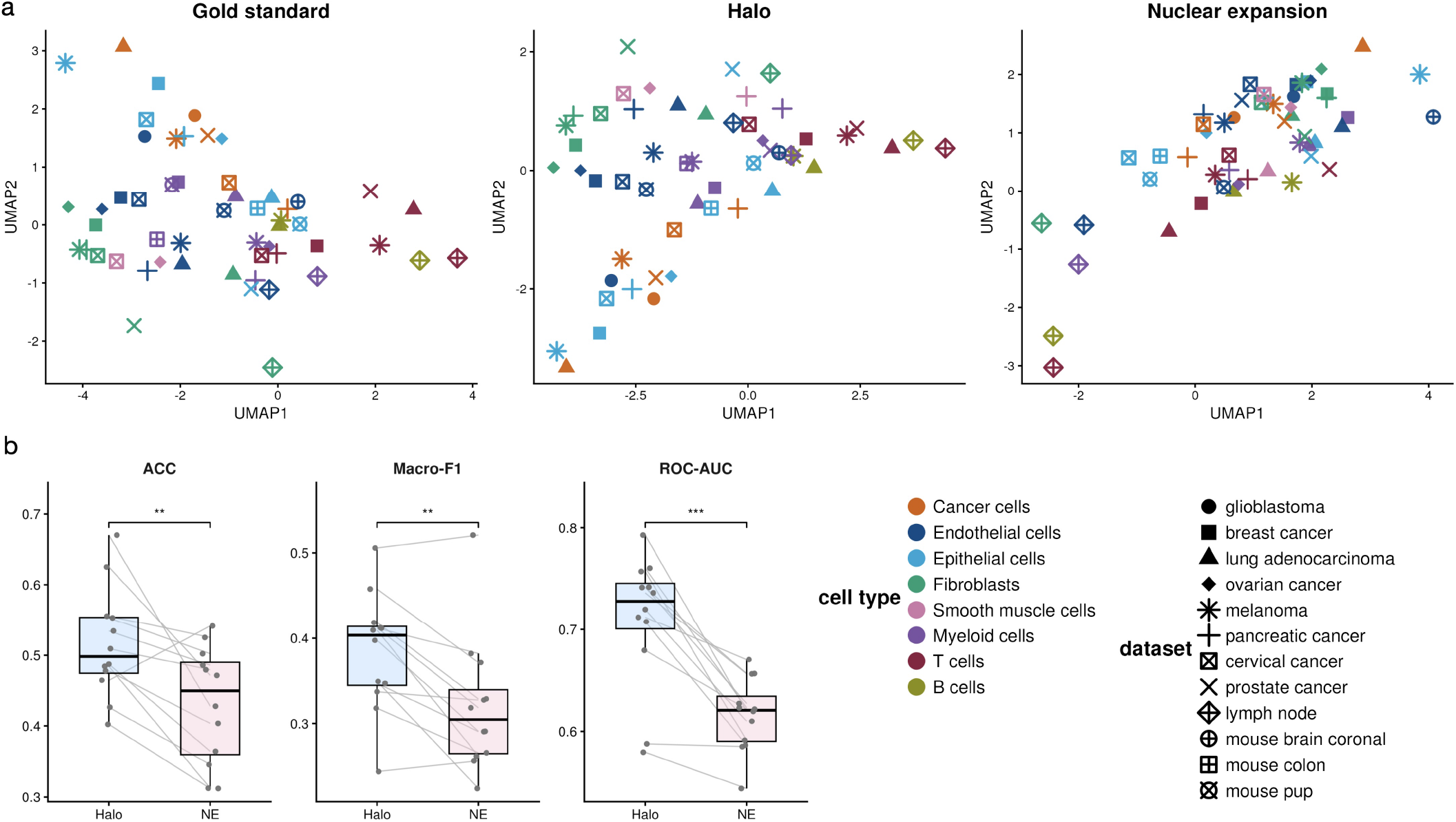
UMAP visualization and random forest analysis of morphological features. **a**, Cell-type centroids across datasets in UMAP space. **b**, Accuracy (left), macro-F1 (middle), and macro ROC-AUC (right) of the random forest classifier. *** indicates p-value < 0.001, and ** indicates p-value < 0.01.

## Discussion

In this study, we introduce Halo, a pretrained model that enables accurate whole-cell segmentation using nuclei staining images and spatial locations of RNA transcripts as input. By converting RNA spatial locations into a pseudoimage, Halo effectively integrates these two data modalities and allows the use of existing segmentation methods developed for imaging data. We demonstrate that Halo substantially improves segmentation accuracy and downstream analyses compared with the current state-of-the-art approach, nuclear expansion.

While Halo currently relies on the Cellpose-SAM model, the training data can be directly used by other models that support cell or instance segmentation. Therefore, Halo can be readily improved as more powerful models pretrained on larger datasets become available in the future. To facilitate such developments, we have made the training data openly accessible.

When constructing pseudoimages from the spatial locations of RNA transcripts, Halo does not distinguish transcripts from different genes. Although transcripts from certain genes may preferentially localize to specific subcellular regions, incorporating such information would require the same gene set to be used during both training and inference on new datasets. Because different ST platforms and experiments often measure different gene panels, Halo does not differentiate transcripts by gene identity, which helps maximize generalizability across datasets with varying gene panels.

## Methods

### Halo

#### Data collection and preprocessing

To train the Halo models, we collected 15 10x Genomics Xenium samples from the 10x Genomics website, spanning 12 distinct human and mouse tissue types. Detailed information for all samples is provided in Supplementary Table 1. For each sample, three types of information were collected: the DAPI nuclei staining image, whole-cell boundaries derived from multimodal staining, and the spatial locations of RNA transcripts. DAPI images were intensity-normalized by capping pixel values at the 99^th^ percentile and then scaling all intensities to the range [0, 1]. For the transcript data, only gene transcripts with a quality score (Q-score) ≥ 20 were retained.

From each sample, we randomly selected 1,000 image tiles, each containing at least 200 pixels within cellular regions and a minimum of 15 cells, yielding a total of at least 180,000 cells across all samples. The tiles from each sample were randomly partitioned into 800 training tiles and 200 test tiles. All training tiles were used to train Halo. For downstream performance evaluation, only the test tiles from 12 of the 15 samples were retained, as gene expression matrices for the remaining three samples could not be reliably retrieved due to technical issues with the 10x Space Ranger pipeline.

#### Transcript-density pseudo-image

Let 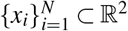 denote the spatial coordinates of the *N* detected RNA transcripts, where 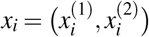 is the two-dimensional location of transcript *i*.

We construct a transcript-density pseudo-image *f* (*u*) by placing a two-dimensional Gaussian kernel at each transcript location and summing their contributions:

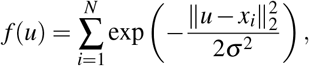

where *u* ∈ ℝ^2^ denotes a spatial coordinate and *σ* = 2.5 controls the spatial spread of each transcript signal.

In practice, *f* (*u*) is evaluated on the discrete pixel grid of the image domain to obtain the transcript-density pseudo-image. To ensure compatibility with the scaled DAPI image, the pixel values of the transcript-density pseudo-image were linearly scaled to the range [0, 1]. The scaled DAPI image and the scaled transcript-density image were then concatenated to form a two-channel input image.

#### Model training

Cellpose-SAM (version 4.0.7) was trained to predict whole-cell boundaries from the two-channel input image. All training tiles across all training samples were included in the training process. The model was trained for 300 epochs using a learning rate of 0.001 and a weight decay of 1e-5.

#### Model inference

The trained Halo model can be applied to new spatial transcriptomics datasets. For each dataset, a transcript-density pseudo-image must first be generated, and the corresponding two-channel input image constructed using the same procedure described above. The resulting two-channel images are then used as input to the trained Halo model for whole-cell boundary prediction.

### Competing method

The nuclear expansion method was included as a baseline for comparison. Specifically, 10x Xenium Ranger was applied to all test datasets in this study using its built-in nuclear expansion strategy, in which nuclear masks were uniformly expanded by 5*µ*m, unless the expansion encountered the boundary of a neighboring cell, to approximate whole-cell boundaries.

### Performance evaluations: Intersection-over-Union (IoU)

#### Image IoU

The image-level IoU was computed separately within each test image tile. Both the ground-truth and predicted whole-cell boundaries were represented as labeled images, where background pixels were assigned a value of 0 and each individual cell was assigned a unique positive integer label.

Let *G*_*i*_ denote the pixel set corresponding to ground-truth cell *i*, and let *P*_*j*_ denote the pixel set corresponding to predicted cell *j*. The intersection-over-union between *G*_*i*_ and *P*_*j*_ was defined as

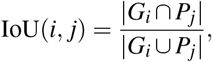

where | · | denotes the number of pixels in the set. This definition produces an IoU matrix whose rows correspond to ground-truth cells and whose columns correspond to predicted cells.

For each predicted cell *j*, its IoU score was defined as

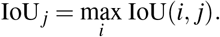

The final image-level IoU was computed as the mean of IoU _*j*_ across all predicted cells, and results were averaged across tiles.

#### Gene IoU

For each cell, transcripts were assigned based on spatial containment within the corresponding segmentation mask. Only transcripts that passed quality control filtering were included in the analysis.

For a given ground-truth cell *i* and predicted cell *j*, let 𝒦_*i*_ and ℒ_*j*_ denote the sets of transcripts assigned to each cell, respectively. The gene-level intersection-over-union was defined as

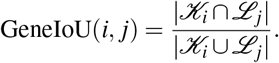

For each predicted cell *j*, its gene IoU score was defined as

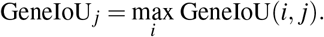

Gene IoU scores were averaged across predicted cells within each tile and subsequently averaged across tiles to obtain dataset-level summary statistics.

### Performance evaluation: cell clustering by gene expression

#### Cell alignment

To facilitate comparison of cells obtained from different methods, the cell boundary masks generated by Halo or by the nuclear expansion strategy were aligned to the ground-truth cell boundary masks. Specifically, a one-to-one correspondence was established between a cell *i*, whose mask was obtained from Halo, and a cell *j* in the ground truth if the following three criteria were satisfied: (1) the cell-level IoU between *i* and *j*, calculated as described above, was at least 0.3; (2) among all cells identified by Halo, cell *i* had the highest IoU with cell *j*; and (3) among all ground-truth cells, cell *j* had the highest IoU with cell *i*.

The same procedure was applied to establish cell alignment between cells identified by the nuclear expansion strategy and the ground truth.

#### Cell clustering

For the gold-standard cell boundaries, as well as those generated by Halo or the nuclear expansion strategy, the corresponding cell-by-gene count matrix was computed using 10x Xenium Ranger. Only cells that were successfully aligned in the previous step were retained for downstream analysis. Each gene expression count matrix was subsequently processed using the Python package `Scanpy`. Briefly, the count matrix was library-size normalized and log-transformed, followed by principal component analysis (PCA). A *k*-nearest neighbor graph was then constructed in the PCA space, and Leiden clustering was performed, with the resolution parameter iteratively tuned to obtain a target of *k* = 10 clusters. If *k* = 10 clusters cannot be achieved through iteration, then *k* = 9 or *k* = 11 is accepted. If neither is attainable, then *k* = 8 or *k* = 12 is considered.

#### Cell type annotation

Cell type annotation was conducted for each cluster using the `sctype` R package, based on curated cell type specific marker genes. For each cancer dataset, a cancer specific marker gene set was additionally applied to identify malignant cell populations. The full list of cell types and their corresponding marker genes used in this study is provided in Supplementary Table S2.

#### Cluster agreement

Agreement between clustering results from Halo or the nuclear expansion strategy and the ground-truth reference was evaluated on aligned cells. Concordance was quantified using the adjusted Rand index (ARI), adjusted mutual information (AMI), homogeneity, and completeness, computed with `adjusted_rand_score`, `adjusted_mutual_info_score`, `homogeneity_score`, and `completeness_score` from the `scikit-learn` library.

### Performance evaluation: cell morphology

#### Morphological features

For each segmented cell, a set of morphological features was computed from its corresponding cell boundary mask.

##### Area

Cell area was defined as the area enclosed by the segmented cell boundary polygon.

##### Perimeter

Cell perimeter was computed as the total length of the segmented polygon boundary.

##### Feret’s diameter

Feret’s diameter was defined as the maximum pairwise Euclidean distance between any two points on the convex hull of the cell polygon.

##### Eccentricity

Cell eccentricity was quantified using PCA of the polygon vertex coordinates. It was defined as 1 − *λ*_2_*/λ*_1_, where *λ*_1_ and *λ*_2_ denote the variances explained by the first and second principal components, respectively. Values closer to 1 indicate more elongated shapes, whereas values closer to 0 indicate more isotropic shapes.

##### Roundness

Roundness was defined as the ratio between the cell area and the area of its minimum bounding circle. Values approaching 1 indicate shapes that closely approximate a circle and efficiently fill their bounding circle, whereas smaller values reflect elongated or irregular morphologies.

##### Circularity

Circularity was computed as 4*πA/P*^2^, where *A* denotes the cell area and *P* denotes the cell perimeter. A perfect circle yields a circularity of 1, whereas lower values indicate increasingly irregular or elongated boundaries.

##### Solidity

Solidity was defined as the ratio between the cell area and the area of its convex hull. This metric quantifies boundary concavity, with values near 1 indicating nearly convex shapes and lower values indicating the presence of indentations or fragmented outlines.

##### Aspect ratio

Aspect ratio was computed from the minimum-area rotated bounding rectangle of each cell as the ratio between the rectangle width and height. Larger values correspond to more elongated shapes, whereas values near 1 indicate more equiaxed cell geometries.

#### UMAP dimension reduction

All morphological features described above were used as input. Each feature was standardized across cells using z-score transformation. Uniform Manifold Approximation and Projection (UMAP) was then performed using the uwot package (version 0.2.3), with n_neighbors = 30, min_dist = 0.01, and Euclidean distance.

To visualize the global organization of cell morphologies, tissue-level centroids were computed by averaging embedding coordinates of cells within each dataset–cell type combination. These centroids were then plotted, with colors indicating cell types and point shapes representing datasets.

#### Random forest classification

To evaluate whether cell morphological features could discriminate cell types, we trained a random forest classifier separately for each tissue. For each tissue, cells labeled as unknown or unassigned were excluded before model fitting. We then restricted the analysis to the five most abundant cell types within that tissue to reduce extreme class sparsity and ensure stable multiclass classification. Morphological features were extracted from the selected feature columns and only numeric features were retained for modeling.

For each tissue-specific dataset, the samples were divided into training and test sets using a stratified split, such that each cell type was represented in both sets while approximately 20% of cells were assigned to the test set. Cell type labels were treated as the response variable, and the morphological features were used as predictors. Random forest models were trained using the ‘randomForest’ R package with class weighting enabled, where class weights were set inversely proportional to class frequency in the training set to partially mitigate class imbalance. Unless otherwise specified, models were trained with 500 trees for tissue-level analyses.

Model performance was evaluated on the held-out test set using three complementary metrics: accuracy, macro-F1, and macro ROC-AUC.

Accuracy was defined as the proportion of correctly classified cells among all cells in the test set:

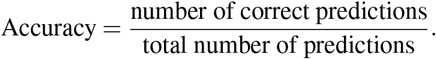

This metric summarizes overall classification performance but may be influenced by class imbalance.

Macro-F1 was used to assess balanced multiclass performance across cell types. For each class, precision and recall were calculated in a one-vs-rest manner:

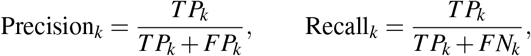

where *T P*_*k*_, *FP*_*k*_, and *FN*_*k*_ denote the true positives, false positives, and false negatives for class *k*, respectively. The class-specific F1 score was then computed as

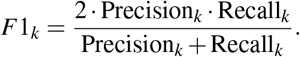

Macro-F1 was obtained by averaging the F1 scores across all classes:

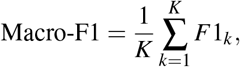

where *K* is the number of classes. This metric gives equal weight to each class and is therefore more sensitive to minority-class performance than accuracy.

Macro ROC-AUC was used to quantify the probability-based discrimination ability of the classifier. For each class, a one-vs-rest receiver operating characteristic (ROC) curve was constructed from the predicted class probabilities, and the corresponding area under the curve (AUC) was computed. The final macro ROC-AUC was defined as the average of the class-specific AUC values:

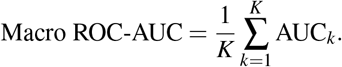

These metrics reflect how well the model separates each of the 5 most predominant cell types from the others.

The same procedure was applied to cell masks obtained from the nuclear expansion strategy and from the ground truth.

## Acknowledgments

The project was supported by the National Institutes of Health under Award Number U54AG075936 and R35GM154865.

## Author contributions

Z.J. conceived the study. X.Z. developed the method and conducted the analysis. X.Z. and Z.J. wrote the manuscript.

## Code availability

The Halo software package and the finetuned model weights are freely available at https://huggingface.co/XYZ1998/ `Halo`, and the training data are publicly available at https://zenodo.org/records/19201256.

